# Structure-Function Relationship in Electrical and Hemodynamic Brain Networks: Insights from EEG and fNIRS during Rest and Task States

**DOI:** 10.1101/2024.04.27.591444

**Authors:** Rosmary Blanco, Maria Giulia Preti, Cemal Koba, Dimitri Van De Ville, Alessandro Crimi

## Abstract

Identifying relationships between structural and functional networks is crucial for understanding the large-scale organization of the human brain. The potential contribution of emerging techniques like functional near-infrared spectroscopy to investigate the structure-functional relationship has yet to be explored. In our study, we characterize global and local structure-function coupling using source-reconstructed Electroencephalography (EEG) and Functional near-infrared spectroscopy (fNIRS) signals in both resting state and motor imagery tasks, as this relationship during task periods remains underexplored. Employing the mathematical framework of graph signal processing, we investigate how this relationship varies across electrical and hemodynamic networks and different brain states. Results show that fNIRS structure-function coupling resembles slower-frequency EEG coupling at rest, with variations across brain states and oscillations. Locally, the relationship is heterogeneous, with greater coupling in the sensory cortex and increased decoupling in the association cortex, following the unimodal to transmodal gradient. Discrepancies between EEG and fNIRS are noted, particularly in the frontoparietal network. Cross-band representations of neural activity revealed lower correspondence between electrical and hemodynamic activity in the transmodal cortex, irrespective of brain state while showing specificity for the somatomotor network during a motor imagery task. Overall, these findings initiate a multimodal comprehension of structure-function relationship and brain organization when using affordable functional brain imaging.

## Introduction

The relationship between brain structure and function is a fundamental concept in neuroscience [1, 2]. Clarifying the interplay or disturbances between structure and function is not only crucial for unveiling the neural mechanisms underlying behaviors, cognition, and disability [3, 4, 5, 6, 7, 8, 9] but also holds significance in terms of early diagnosis and guiding therapeutic interventions [10]. Contemporary imaging technology enables the high-throughput reconstruction of neural circuits across various spatiotemporal scales. Anatomically, diffusion magnetic resonance imaging (dMRI) and diffusion tensor imaging (DTI) visualize physical connections and reconstruct nerve fibers, respectively [11, 12, 13]. Functionally, techniques like functional magnetic resonance imaging (fMRI) and functional near-infrared spectroscopy (fNIRS) detect changes in hemoglobin oxygenation/deoxygenation, serving as indirect indicators of neuronal activity with high spatial resolution [14, 15, 16]. Magnetoencephalography (MEG) and electroencephalography (EEG), instead, measure magnetic and electrical activity generated by groups of neurons with high temporal resolution, respectively [17, 18, 19, 20, 21]. The relationship between functional activity and the underlying structural connectome is usually evaluated using whole-brain diffusion MRI-derived structural connectivity and functional connectivity at a desired temporal scale (either neural or hemodynamical). However, although the neurophysiological and the hemodynamic activity attempt to capture the same underlying biological process, they are sensitive to different physiological mechanisms at different timescales [1, 22, 23] with hemodynamic activity indirectly reflecting the underlying patterns of electrical neural activity due to slow neurovascular coupling [24, 25]. Evidence indicates that the structure-function relationship is not uniform across brain regions, pointing to a more robust coupling in the unimodal cortex, which is sensitive to immediate changes in the sensory environment, and a weaker association in the transmodal cortex, which is sensitive to a prior context [26, 27, 28, 29, 30, 31]. It has been suggested that this regional heterogeneity may potentially reflect underlying molecular and cytoarchitectural gradients, highlighting a hierarchy of time scales of intrinsic fluctuations across the cortex [2, 32, 33, 34, 35, 36]. Moreover, it was reported that the structure–function relationship is modality-dependent, suggesting that functional connections estimated from longer time windows overlap significantly with the underlying structural connections. However, a structural-functional mismatch may arise with shorter time windows due to distributed delays between neuronal populations, leading to transient phase (de-)synchronization [37, 38, 39]. This raises the possibility that the alignment between the relatively slower functional activity captured by the hemodynamic network and the faster functional activity captured by the electrical network may systematically vary across the cortex. Evidence suggests that the functional architectures of EEG/MEG and fMRI are consistent early in the cortical hierarchy, presumably reflecting activity related to instantaneous changes in the external environment. Conversely, as we move up the hierarchy, there is a gradual separation between the two architectures, suggesting that they are differently modulated by endogenous inputs and contextual information [28, 40, 41]. However, it remains unclear how functional networks at different time scales relate to one another and their common structural substrate.

Moreover, previous studies have primarily focused on resting-state activity as inferred from EEG, MEG, and fMRI data, at both global and local levels [42, 43, 44, 45, 46, 46, 47, 48, 49]. However, the extent to which this relationship varies across resting and task-specific periods has not been fully explored. Additionally, while fNIRS has recently demonstrated promise in probing resting-state functional connectivity [41, 50, 51, 52, 53, 54], its potential contribution to investigating the structure-functional relationship has yet to be explored. This recent neuroimaging technique offers improved cost-effectiveness and portability compared to fMRI. Additionally, fNIRS provides a high temporal resolution, typically ranging from 2 to 10 Hz [53, 55], in contrast to the typical fMRI whole-brain scan rate of 0.5 Hz.

In our study, we leverage the graph signal processing (GSP) framework and the harmonic modes theory to map large-scale cortical dynamics in source-projected EEG and fNIRS data. We aimed to explore the hypothesis that the relationship between structural connectivity and functional patterns might vary between electrical and hemodynamic networks, given their different sensitivity to various physiological mechanisms at different timescales [1, 22, 23]. Additionally, we sought to investigate how this relationship varies across cognitive states, which might reflect underlying organizational principles governing behavior. To our knowledge, this study represents the first investigation into the structure-functional relationship using fNIRS data and comparing it with EEG.

The introduction of GSP has offered a novel framework for combining structure-function analyses, allowing the extraction of harmonic basis functions from the structural connectivity (SC), which then serves to obtain a graph-spectral representation of the data [26, 56, 57, 58, 59, 60]. In this context, Preti and Van De Ville (2019) introduced the structural-decoupling index (SDI) [26], which quantifies the degree of structure-function dependency for each brain region. We use the SDI to quantify the global (cross-regional) and local (region-wise) (dis)alignment between the structural and functional networks within and between each modality in both resting state (RS) and task. Moreover, we investigate the degree to which regional patterns of structure-function coupling align with the canonical intrinsic functional networks (RSNs) to gain insights into how the underlying anatomy influences functional networks and supports various cognitive functions. This approach involves comparing hemodynamic and electrical activity in a common frame of reference which might shed light on the interaction between neural signaling and oxygenation demands during different brain states. The relevance of understanding the modality dependency of structure-function relationships might have implications for clinical applications, particularly in addressing questions such as identifying the most sensitive modality for detecting changes in functional networks in the presence of disease-specific structural network damage.

## Methods

### Data preprocessing and source space signal reconstruction

The functional data used in this study were acquired from 18 healthy subjects (28.5 ± 3.7 years) through an open dataset [61]. The dataset included synchronous EEG and fNIRS recordings during 1-minute resting state (RS) sessions and 30 trials of 10-second left and right-hand motor imagery tasks (MI) for each participant. EEG data were recorded with 32 electrodes placed according to the international 10-5 system (Fig. 1 (a)) at a 1000 Hz sampling rate (down-sampled to 200 Hz). fNIRS data were collected by 36 channels (14 sources and 16 detectors with an inter-optode distance of 30 mm) (Fig. 1 (b)), following the standardized 10-20 EEG system, at a 12.5 Hz sampling rate (down-sampled to 10 Hz). Two wavelengths at 760 nm and 850 nm were used to measure the changes in oxygenation levels.

**Figure 1.**
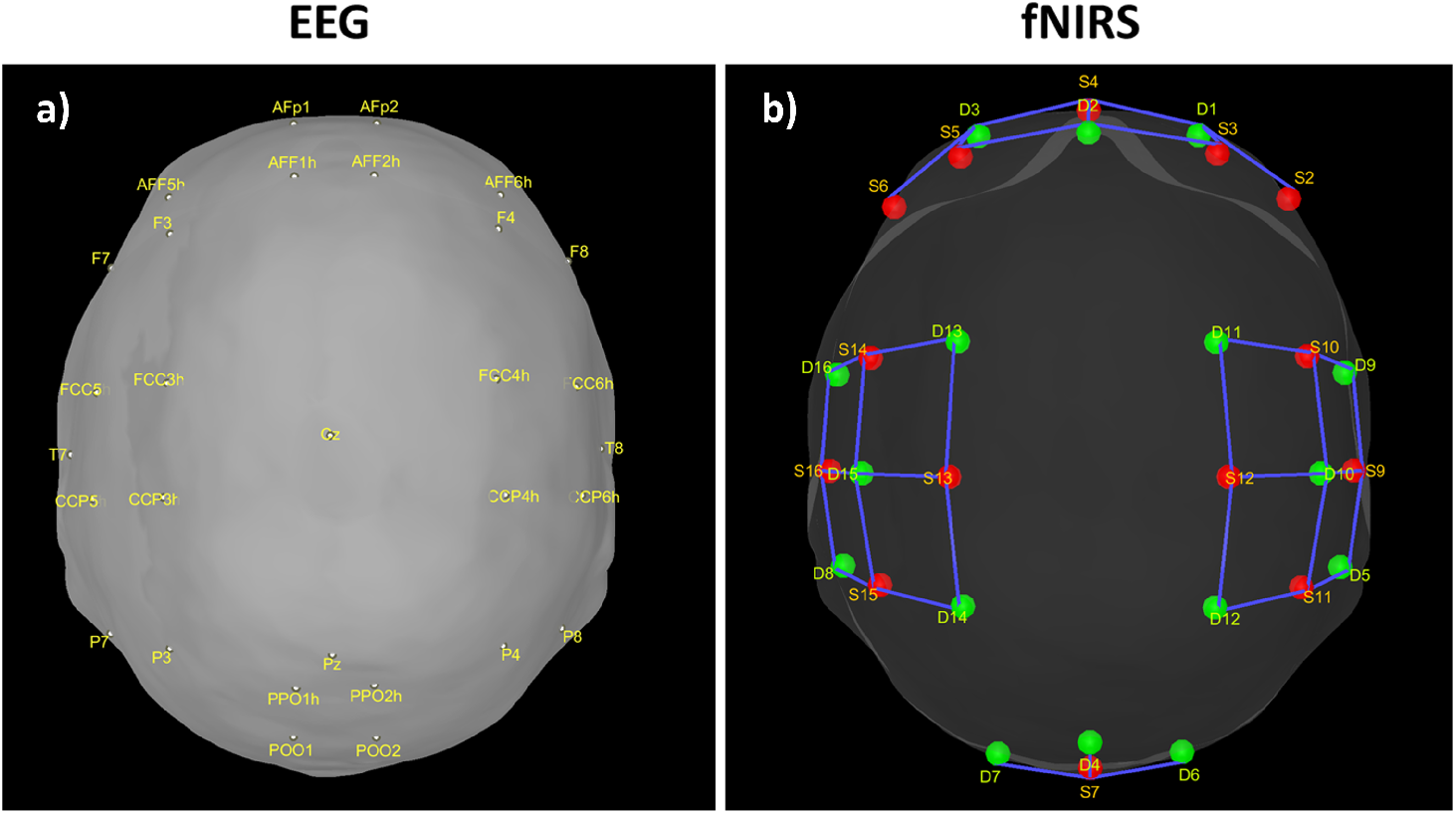
**(a):** EEG electrode locations. **(b):** NIRS optode locations. The red dots are the sources and the green dots are the detectors.

### Resting-state and task functional fNIRS source timeseries

Resting-state and task fNIRS scans were processed using the MNE toolbox [62] and Brainstorm software [63]. Optical density (OD) transformation was applied to the signals, and the scalp-coupled index (SCI) was used to assess signal quality.

Subjects with more than 50% of channels displaying an SCI *<* 0.7 were excluded. The signals were bandpass filtered with cutoff frequencies of 0.02-0.08 Hz using a finite impulse response (FIR) filter. Time segments with excessive head movements, identified through the global variance in temporal derivative (GVTD) metric [64], were rejected. To address systemic physiological effects, principal component analysis (PCA) was applied, and components with the highest spatial uniformity value, indicative of superficial skin responses [65], were removed. Task-related time epochs from 0 to 10 seconds relative to the stimulus onset were averaged across trials for each subject during both left and right MI. The diffuse optical tomography (DOT) method was implemented to reconstruct the signals in the source space for both the RS and task-related time epochs. A five-tissue segmentation of the Colin27 brain template was used to compute the forward model (sensitivity matrix). The fluences of light for each optode were estimated using Monte Carlo simulations with a number of photons equal to 10^8^ and projecting the sensitivity values within each vertex of the cortex mesh. The forward model was computed from fluences by projecting the sensitivity values within each vertex of the cortex mesh using the Voronoi-based method [66]. The depth-weighted minimum norm estimate (depth-weighted MNE) method was utilized to estimate the sources. Parcellated time series were then estimated and mapped onto the three-dimensional space using a reduced set of 44 regions of interest (ROIs) from the Desikan-Killiany atlas instead of 82, due to the incomplete coverage of optodes across the scalp. The source time series were then converted into oxygenated hemoglobin (HbO) and deoxygenated hemoglobin (HbR) by applying the modified Beer-Lambert transformation.

### Resting-state and task functional EEG source timeseries

Resting-state and task EEG scans underwent processing using the MNE toolbox [62] and Brainstorm software [63]. The preprocessing pipeline involved re-referencing with a common average reference and applying a second-order zero-phase Butterworth type FIR high-pass filter with a cutoff of 1 Hz. Independent component analysis (ICA) was performed using the Infomax algorithm for artifact removal. Artifacts were identified and eliminated through visual inspection of various diagnostic measures for each independent component (IC). Subsequently, the cleaned data were low-pass filtered (second-order zero-phase Butterworth type FIR filter) with a cutoff of 45 Hz. Task-related time epochs from 0 to 1 second relative to the stimulus onset were averaged across trials for each subject during left and right MI. Source estimation (ESI) for both the RS and task-related time epochs was performed using the standardized Low-Resolution brain Electromagnetic Tomography (sLORETA) method on an MRI template (Colin27). Data covariance regularization was applied using the median eigenvalue method to address variable source depth effects. Source orientations were constrained as normal to the 15000-vertex Colin27 surface, and parcellated time series were then estimated and mapped in the same 3D space of the Desikan-Killiany atlas as fNIRS for comparison. The source time series were then decomposed into the typical oscillatory activity by band-pass filtering: *delta* (1 *−* 4*Hz*), *theta* (4 *−* 7*Hz*), *alpha* (8 *−* 15*Hz*), *beta* (15 *−* 25*Hz*), and *gamma* (25 *−* 45*Hz*).

### Structural and diffusion MRI

The group-consensus structural connectome was computed using the structural connectivity matrices derived from the CONNECT/ARCHI database. The dataset consisted of structural connectivity matrices and resting-state functional data (fMRI) obtained from 78 subjects in the ARCHI database. Individual structural connectivity matrices were constructed from diffusion MRI, with connection strength determined by fibers reconstructed through a tractography algorithm [67, 68, 69]. The connectivity data were then mapped onto the Desikan atlas and subsequently reduced to 44 ROIs to align with the functional data. The group-consensus structural connectome, representing the entire population, was derived by averaging the structural connectivity (SC) values across subjects.

### Structure-Function Coupling Features

As an indicator of the structure-function relationship, the SDI was computed using the Graph Signal Processing (GSP) framework detailed in [26]. This analysis was performed for each subject, modality (EEG and fNIRS), and condition (resting state (RS), right and left motor imagery (MI) tasks). GSP provides a method for analyzing and representing graph signals in a spectral domain, utilizing the eigenvectors of the Laplacian matrix as fundamental basis functions. The core idea is that these fundamental basis functions can serve as ‘building blocks’ for brain activity within the same domain. Each regional brain activity measurement *s*(*t*) at different time points *t* is considered as a graph signal, with each brain region represented as a node within the structural connectivity (SC) graph, described by the adjacency matrix **W**. The building blocks, referred to as harmonics, are obtained through an eigendecomposition of the symmetrically normalized Laplacian matrix **L** of **W**, denoted as:

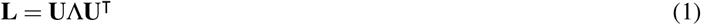

where **U** = [*µ*_1_, *µ*_2_, …, *µ*_*N*_], is the matrix of eigenvectors *µ*_*i*_, and Λ = diag(*λ*_1_, *λ*_2_, …, *λ*_*N*_) is the diagonal matrix of eigenvalues *λ*_*i*_. Each eigenvalue *λ*_*i*_ can be interpreted as the spatial frequency of the corresponding structural harmonic (eigenvector) *µ*_*i*_.

Subsequently, the graph Fourier transform (GFT) is utilized on the graph signal s*(t)* to extract their graph Fourier coefficients *ŝ*. Given the parcellated source reconstructed signal as an array of dimensionality *N × T*, where *N* is the number of brain regions (*N* = 44) and *T* is the number of time points, each *N* × 1 column of this array is an activation pattern *s*(*t*) indexed by time *t* and their graph Fourier coefficients *ŝ* represented as:

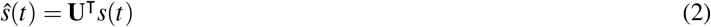

where **U**^T^ is the transposed of the eigenvector matrix **U**. Then, the filtering process, in the graph spectral domain, is applied to extract relevant frequency components, by using ideal low-pass (**F**_low_) and high-pass (**F**_high_) filters. These filters are represented as diagonal matrices with non-zero entries corresponding to the components to be retained. The cut-off frequency is determined based on a median split criterion, which ensures that the energy distribution in the retained frequency components is balanced between the lower and the upper portion of the signal spectrum. For each subject, EEG band, oxy- deoxyhemoglobin, and conditions (RS, left and right MI) we computed the power spectral density (PSD) using the Matlab *pwelch* function and identified the cut-off frequency that divided the averaged spectrum into two portions of approximately equal energy. The filtered graph signals are transformed back into the temporal domain by applying the inverse graph Fourier transform. This results in a low-frequency functional activity component *s*(*t*)_*c*_ = **UF**_low_**U**^*T*^ *s*(*t*), which is coupled to the structure, and a high-frequency functional activity component *s*(*t*)_*d*_ = **UF**_high_**U**^*T*^ *s*(*t*), more decoupled from the structure. The ratio between these two signal portions yields the SDI, quantified as:

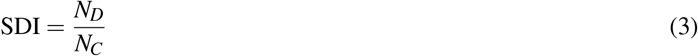

where *N*_*C*_ (coupling) and *N*_*D*_ (decoupling) are the filtered spectral coefficients computed as the *l*2-norm across time of the coupled *s*(*t*)_*c*_ and decoupled *s*(*t*)_*d*_ portions for each brain region. The SDI map for each subject is obtained for both the RS and the averaged SDI values of the left and right MI task (named task) for each EEG frequency band and fNIRS hemoglobin type.

### Statistical analyses

We seek to elucidate how structural-function coupling systematically varies within and between modalities (EEG and fNIRS), globally across regions of interest (ROIs) and over the neocortex (for each ROI), from the perspective of topological, intrinsic functional organization during RS and task conditions. The statistical analysis was performed using the Matlab toolbox (version 2020a) [70]. We assessed the normality of data distribution (SDI across subjects and frequency bands for EEG and across subjects and hemoglobin types for fNIRS) by using the *Skewness* Matlab function. Given that the data did not exhibit a normal distribution (EEG: RS Skewness = -0.6652, Task Skewness = -0.6393; fNIRS: RS Skewness = -0.5493, Task Skewness = -0.5107), we opted for a non-parametric statistical tests.

### Global structure-function coupling

Our initial hypothesis aimed to determine significant differences in the overall distribution of SDI values between modalities for both RS and task conditions. To test this we used the Wilcoxon signed-rank test:

1. to compare EEG SDI values across ROIs, subjects, and frequency bands with fNIRS SDI values across ROIs, subjects, and hemoglobin types (HbO, HbR) for each condition.
2. to compare the SDI values within each modality for each condition. Specifically, we compared SDI values between each pair of frequency bands for EEG and between HbO and HbR for fNIRS for both RS and task.
3. to examine the role of different oscillations and hemoglobin types in modulating the dynamics of the structure-function relationship in response to the transition from RS to task. we compared SDI values within each modality between conditions, assessing each pair of frequency bands and HbO with HbR.
4. to compare the SDI values between modalities for each condition. Specifically, we compared each pair of EEG frequency bands with each HbO and HbR fNIRS.

False discovery rate (FDR) correction was applied to account for multiple comparisons, controlling the family-wise error rate at 0.05. Pearson’s correlation coefficient (*r*) was used to assess the similarity of SDI values across subjects and ROIs within (comparing the frequency bands and the hemoglobin types) and between modalities (comparing each frequency band with each hemoglobin type).

### Local structure-function coupling

Our second hypothesis aimed to determine how regional heterogeneity influences the structure-function relationships between modalities for both conditions. Regional variations in SDI values across subjects were used to determine the spatial distribution of structure-function coupling for each modality and condition. Then, to identify brain areas where the coupling/decoupling patterns differ between modalities, the Wilcoxon signed-rank test was employed to compare the regional SDI of each EEG band with HbO and HbR separately for both RS and task. False discovery rate (FDR) correction was applied to account for multiple comparisons, controlling the family-wise error rate at 0.05.

To shed light on how the underlying anatomy influences functional networks and supports various cognitive functions, we investigated how regional patterns of structure-function coupling align with the canonical intrinsic functional networks. We categorized regions based on their association with macro-scale intrinsic networks [71] and computed the average SDI values across all regions (defined by the Desikan atlas) belonging to each of the five functional networks: the default mode network (DMN), the attentional network (AN) which includes both dorsal (DAN) and ventral (VAN) attention network, the frontoparietal network (FPN), the visual network (VIS), and the somatomotor network (SMN) [72, 73]. Subsequently, we performed comparisons between each EEG-band network and their corresponding HbO and HbR network separately, employing the Wilcoxon signed-rank test for both RS and task conditions. Additionally, we conducted comparisons between cross-band EEG network and its corresponding HbO and HbR pairs separately. We concatenated the SDI values across frequency bands, averaged all regions belonging to each of the five functional networks, and compared them to the corresponding HbO and HbR pair separately using the Kruskal-Wallis test for both conditions.

## Results

### Global modality-independent structure-function relationship

At the global level, our analysis revealed a notable overall difference in structure-function coupling between EEG and fNIRS for the RS, while no disparity was shown for the task condition (Fig. 2 (a)). This implies that the modalities exhibit more similar patterns of structure-function coupling/decoupling when the brain is engaged in a specific cognitive task. When investigating how the structure-function relationship changes across different EEG frequency bands and oxy- and deoxyhemoglobin in both conditions, we observed a general trend of increased coupling for both modalities during the task period, in contrast to the RS (Fig. 2 (g)), alongside distinct patterns of coupling and decoupling. Specifically, in RS, the alpha oscillations demonstrated stronger coupling with the underlying structure than all other bands, which aligns with the notion that alpha oscillations are linked to an idling or inhibitory state, reflecting a coordinated and synchronized neural network at rest. However, the delta and theta bands exhibited higher coupling than the beta and gamma bands, which aligns with the well-established role of these oscillations in fundamental brain functions, such as the overall integration of neural processes. Interestingly, no significant differences in decoupling patterns were observed between the beta and gamma bands, suggesting a similar degree of structural-functional interaction for higher frequency in the absence of cognitive demand (Fig. 2 (b)).

**Figure 2.**
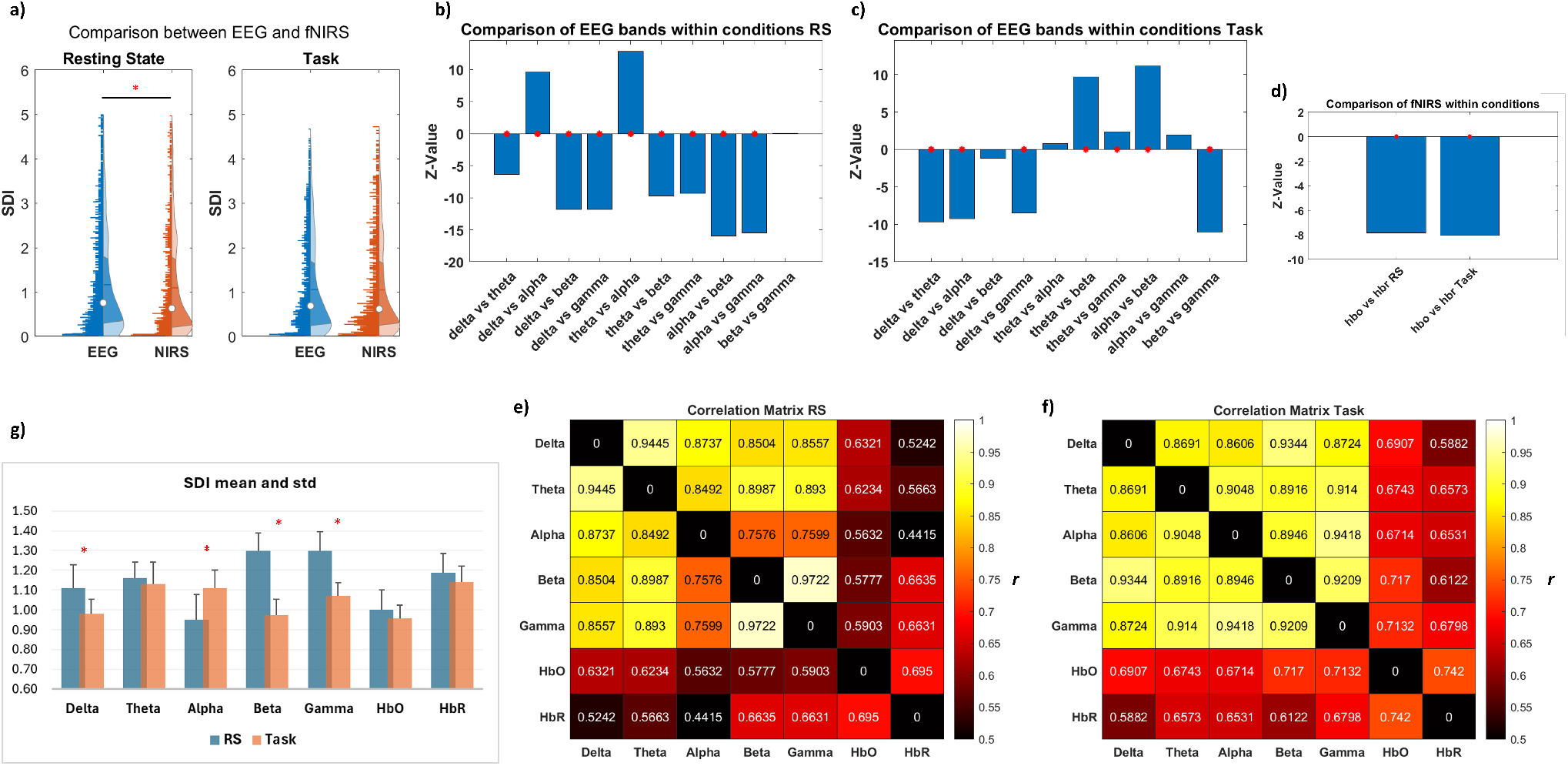
Global structure-function coupling. **(a)**: Overall assessment of SDI differences between EEG and fNIRS across ROIs, subjects, frequency bands, and hemoglobin types, in both RS and task conditions. The red asterisk highlights statistical significance (*p <* 0.05). **(b - c)**: Bar plot illustrating the comparison of SDI between EEG bands, and **(d)**: between HbO and HbR, during RS and task, represented as z-values. A higher absolute z-value indicates a more pronounced difference between the two groups. A positive z-value suggests that the first group exhibits higher values than the second group (indicating more decoupling), while a negative z-value implies lower values in the first group (indicating more coupling). The red asterisk denotes statistical significance (*p <* 0.05). **(e - f)**: Evaluation of similarity within and between modalities (across each EEG frequency band and between HbO and HbR for fNIRS, as well as between each band and each hemoglobin) using Pearson’s correlation coefficient (*r*). **(g)**: mean and std of SDI values, across ROIs and subjects, for each EEG band and fNIRS chromophore in RS and task. The red asterisk denotes statistical significance (*p <* 0.05).

During the task, distinct patterns emerged, shedding light on the interaction between structural and functional dynamics during a cognitive engagement. A stronger coupling in the beta compared to all other bands, might reflect a task-specific modulation of neural interactions, with beta oscillations playing a more prominent role in motor imagery-related processes. Delta band exhibited a higher degree of coupling compared to theta, alpha, and gamma bands, potentially reflecting the involvement of lower oscillations in motor planning and execution. On the contrary, theta and alpha oscillations displayed increased decoupling patterns compared to delta, beta, and gamma bands, suggesting a reduction in the tight relationship between structural and functional profiles during the motor imagery-related task. This may be related to the suppression of alpha activity associated with increased cognitive demands and motor planning (Fig. 2 (c)). For fNIRS, during both conditions, HbO exhibited higher coupling than HbR (Fig. 2 (d)), suggesting that the relationship between structural and functional dynamics is closely linked to the demand for oxygenation.

When examining the similarities between EEG bands, frequency-wise correlations revealed higher similarities between delta and theta bands (*r* = 0.94, *p <* 0.001) and between beta and gamma bands (*r* = 0.97, *p <* 0.001) during resting state (RS), whereas alpha bands exhibited the lowest similarities (*r* = 0.75, *p <* 0.001). This suggests potentially distinct functional dynamics within the alpha range during rest. Conversely, in the task condition, the highest similarities were found between alpha and gamma bands (*r* = 0.94, *p <* 0.001) and delta and beta bands (*r* = 0.93, *p <* 0.001), indicating a reconfiguration of synchronization patterns and coupling mechanisms during the cognitive task. The similarities between HbO and HbR yielded correlation values of *r* = 0.69 (*p <* 0.001) for RS and *r* = 0.74 (*p <* 0.001) for the task (Fig. 2 (e-f))). To give more insight into the role of the different oscillations and hemoglobins in modulating the dynamics of the structure-function relationship in response to the transition from RS to task, we compared each EEG frequency band and each oxy- and deoxyhemoglobin fNIRS between conditions. The observed patterns indicate that different EEG frequency bands exhibit varied dynamics of coupling/decoupling in the transition from RS to MI task. Specifically, higher coupling was observed within the alpha band during the RS accompanied by decoupling trends during the Task. Conversely, higher decoupling characterized the delta, beta, and gamma bands in RS, a trend that shifted towards higher coupling during the task. No notable differences were observed in the theta band. These observations suggest a task-specific modulation of neural activity, possibly indicating the engagement of the faster oscillations in MI tasks, alongside the increased functional demands in this cognitive state (Fig. 3 (a-b)). On the contrary, no significant differences in the structure-function relationship were observed between RS and task for both HbO and HbR, suggesting an overall relatively stable relationship between oxygenated/deoxygenated hemoglobin levels and functional activity across both brain states (Fig. 3 (c-d)).

**Figure 3.**
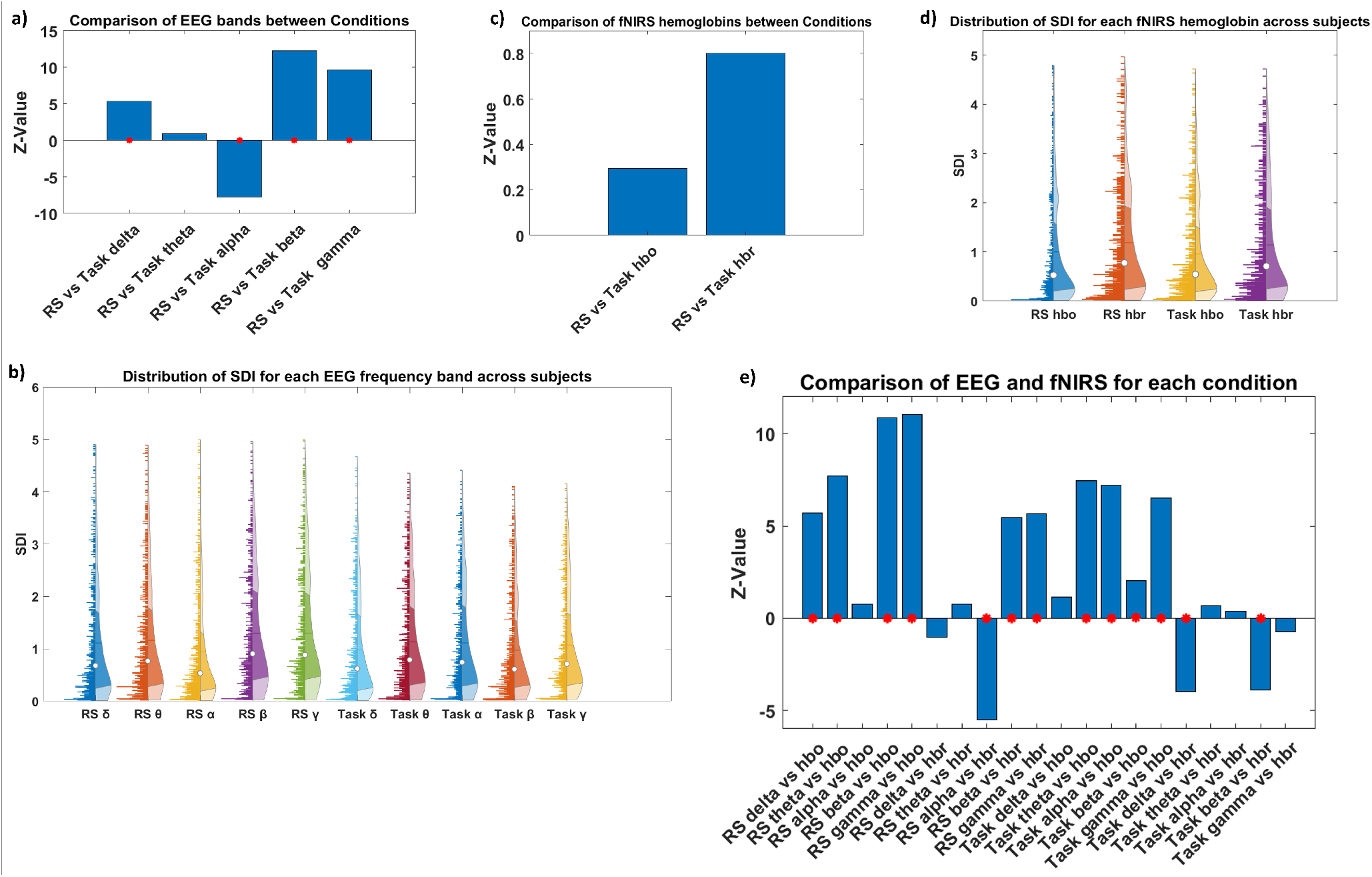
Global structure-function coupling: Bar plot comparing SDI values for each EEG band in **(a)** and for each HbO and HbR in **(c)** across conditions, displayed as z-values. A higher absolute z-value indicates a more significant difference between the two groups. A positive z-value suggests that the first group has higher values than the second group (indicating more decoupling), while a negative z-value implies lower values in the first group (indicating more coupling). The red asterisks indicate statistical significance (*p <* 0.05). **(b - d)**: Illustration of SDI distribution across ROIs and subjects for each EEG band and each hemoglobin for fNIRS, respectively, in both resting state and task conditions. **(e)**: Comparison between modalities (each band with each hemoglobin) for both resting state and task, represented as z-values.

### Global modality-dependent structure-function relationship

Finally, we aimed to explore the interaction between the two neuroimaging modalities by comparing each EEG frequency band and fNIRS hemoglobin measures in both the RS and task conditions. We observed nuanced structure-functional patterns in limited-band EEG compared to those in oxy- and deoxyhemoglobin in the two conditions. Specifically, HbO was more aligned with the alpha band, as no differences were highlighted between them, and demonstrated a higher coupling with the underlying structure compared to the other bands. This effect was particularly pronounced when compared to the faster bands (beta and gamma) during the resting state (RS). Contrarily, HbR more closely resembles the slower frequency bands (delta and theta) while showing higher decoupling than the alpha band, indicating the distinct roles of the two hemoglobin types in relation to electrical activity. During the task, HbO exhibited alignment with the delta band, demonstrating stronger coupling compared to all other rhythms, with less pronounced differences observed with the beta band. Conversely, HbR exhibited higher decoupling than the delta and beta bands while being more aligned with theta, alpha, and gamma oscillations (Fig. 3 (e)).

When examining the similarities between EEG bands with oxy- and deoxyhemoglobin, a higher correlation between hemodynamic (HbO) and slower frequencies (delta *r* = 0.63, *p <* 0.001 and theta *r* = 0.62, *p <* 0.001) was displayed during the RS, and between HbO and faster frequencies (beta and gamma *r* = 0.663, *p <* 0.001) during the tasks. In contrast, HbR shows a higher correlation with faster frequencies in both RS (beta and gamma *r* = 0.71, *p <* 0.001) and Task (gamma *r* = 0.68, *p <* 0.001) conditions (Fig. 2 (e-f))). The summary of statistical tests is reported in Table 1 (a - b) of the Supplementary Materials.

### Local modality-independent structure-function relationship

Our second hypothesis aimed to determine how regional heterogeneity influences the structure-function relationships between modalities and how this heterogeneous relationship evolves between RS and task conditions. We observed a similar spatial distribution of the structure-function patterns across different frequency bands and for both the neuroimaging modalities and conditions. This pattern was characterized by higher coupling in the sensory cortex and increased decoupling in the association cortex. Specifically, robust coupling (lower SDI) is evident in sensorimotor and visual areas, while robust decoupling (higher SDI) is observed in prefrontal and posterior areas, including superior and inferior parietal, and occipital association areas. However, we observe slightly divergent patterns across bands and hemoglobin types. In the RS, the alpha band demonstrates higher coupling in the right prefrontal and the right posterior parietal cortices compared to other frequency bands and hemoglobin types. Slower rhythms (delta and theta) exhibit increased coupling in the right prefrontal area, with HbO showing a similar trend. While faster rhythms (beta and gamma) display more decoupling in the right prefrontal area, with HbR following a similar pattern. During the task, there is a trend of increased coupling across the delta, beta, and gamma bands, as well as with HbO, in the right parietal regions compared to the RS. In contrast, an opposite pattern emerges for alpha oscillation and HbR, showing higher decoupling in the same regions. (Fig. 4). This suggests that engaging in a cognitive task led to a more convergent relationship between neural activity at specific frequency bands, hemodynamic activity (oxygenated hemoglobin), and underlying brain structure in regions associated with higher cognitive functions, reflecting a shift in functional organization to support task-specific processing.

**Figure 4.**
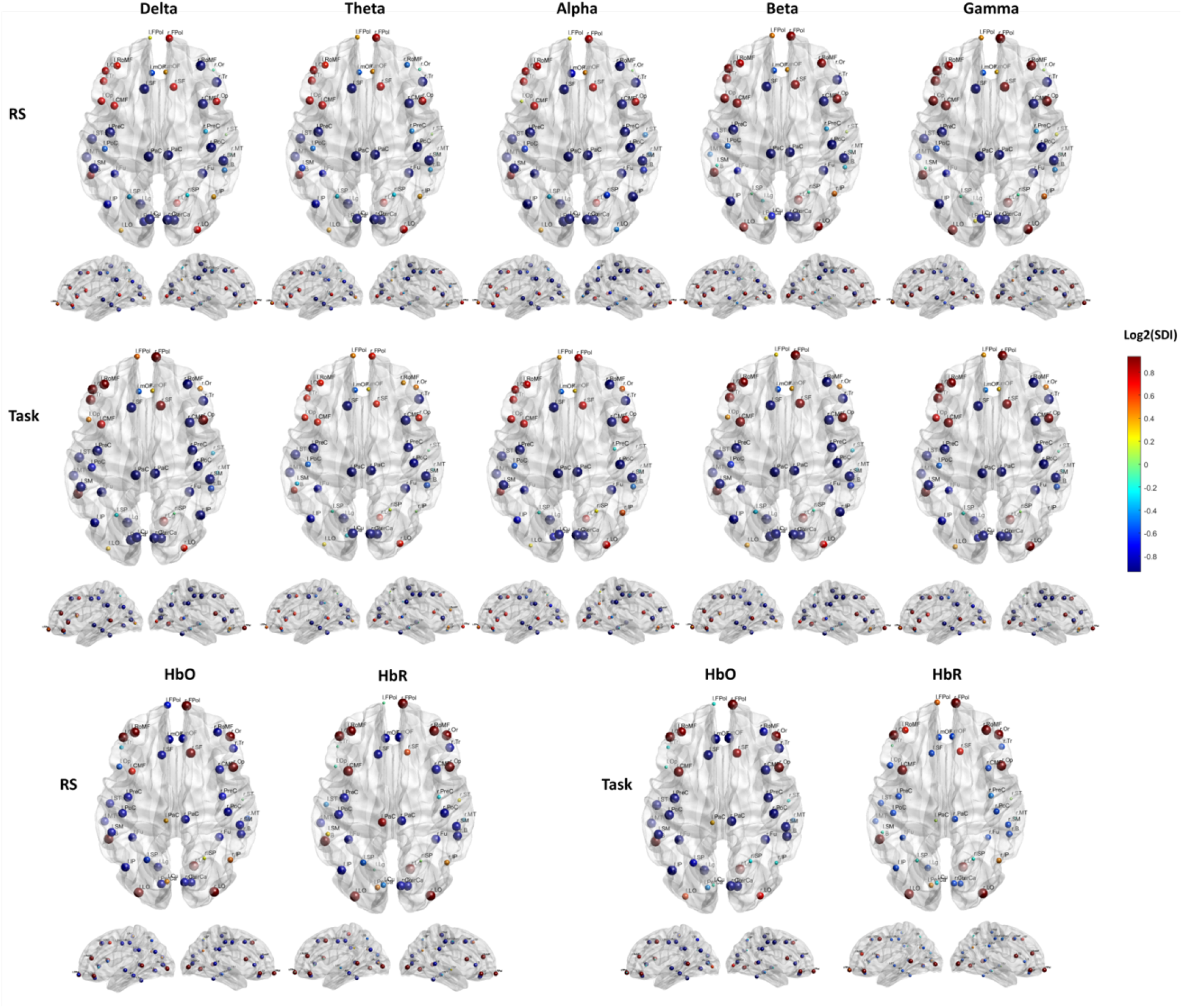
Local structure-function coupling: Spatial distribution of the structure-function coupling across frequency bands (delta, theta, alpha, beta, and gamma) and hemoglobin (HbO and HbR), in RS and task. The blue dots highlight regions more coupled (lower SDI) while the red dots the regions more decoupled (higher SDI).

### Local modality-dependent structure-function relationship

We then quantitatively assessed the degree of alignment between structural connectivity and electrical activity versus hemodynamic activity to discern whether different modalities exhibit similar or divergent patterns of regional structure-function coupling for both RS and task conditions.

We found that the correspondence between HbO and HbR with the EEG bands converged in the profile of the regional structure-function coupling. This convergence manifested as consistent patterns of coupling or decoupling within the same ROI, albeit varying in intensity, indicating a predominant coupling of HbO over the different EEG bands than HbR (Fig. 5 (a)). This discrepancy differed between the RS and task conditions, highlighting a higher level of alignment during the task between HbO and EEG, more pronounced for higher frequencies (beta and gamma), as evidenced by fewer ROIs displaying significant disparities (Table 2 of the Supplementary Materials). Conversely, differences between RS and task conditions were also noted for HbR, but they appeared to be less prominent (a similar number of ROIs displayed significant disparities in both conditions). These differences were marked by a tendency towards greater coupling for electrical activity, particularly evident in the lower frequencies (delta, theta, and alpha) during RS, while across all bands during the Task (Fig. 5 (b)).

**Figure 5.**
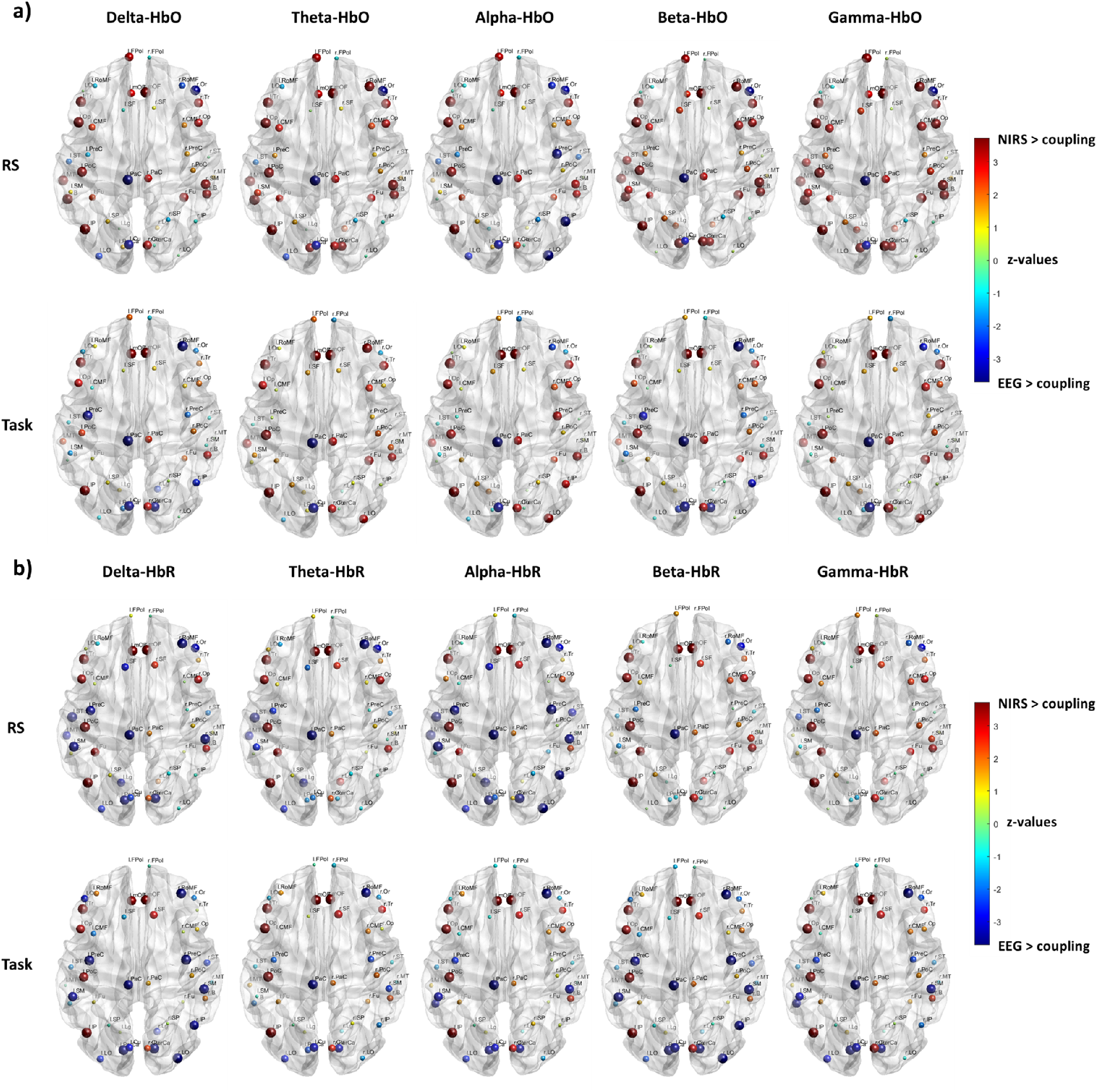
Local structure-function coupling: Spatial distribution of the cross-modal differences in structure-function coupling between **(a)** EEG and HbO, and **(b)** EEG and HbR, in RS and task conditions, represented as z-values. A higher absolute z-value indicates a more significant difference between the two groups, with dot size indicating the absolute z-value. A positive z-value indicates higher coupling for fNIRS, while a negative z-value indicates higher coupling for EEG. Only significant regions of interest (ROIs) are shown (*p <* 0.05), FDR corrected.

Overall, these findings suggest that the spatial organization of EEG structure-function coupling shares similarities with fNIRS but also displays unique characteristics, varying depending on the specific band and condition and indicating differential sensitivity of the modalities to underlying neural processes. For instance, differences in key regions, specifically related to motor imagery tasks, can be observed when comparing electrical and hemodynamic activity. Higher coupling in the dorsolateral prefrontal cortex, premotor cortex, posterior parietal regions, and visual areas is emphasized for EEG over fNIRS in the alpha band during the resting state and in delta and beta bands during the task, while across all frequencies for EEG-HbR.

### Network modality-independent structure-function relationship

To further explore how this distinctive pattern of structure-function coupling observed between electrical and hemodynamic activity is related to functional networks, we categorized the ROIs based on their association with macro-scale intrinsic networks. We observed higher structure-function coupling in the VIS and SMN networks (unimodal cortex) and lower structure-function coupling in the DMN and AN (transmodal cortex) across all EEG frequency bands and fNIRS hemoglobin types. In contrast, the extent of structure-function coupling in the FPN varied depending on the frequency bands, as well as on the brain states, exhibiting a pattern of higher coupling in lower frequency bands (delta, theta, and alpha) and higher decoupling in faster rhythms (beta and gamma) during the RS and an inverted pattern during the task. On the contrary, similar profiles of structure-function coupling in the FPN were observed for oxy- and deoxyhemoglobin between conditions, characterized by a more coupled pattern for HbO and a more decoupled pattern for HbR (Fig. 6) with a general trend toward increased coupling during the task. The summary of statistics is reported in Table 3 of the Supplementary Materials.

**Figure 6.**
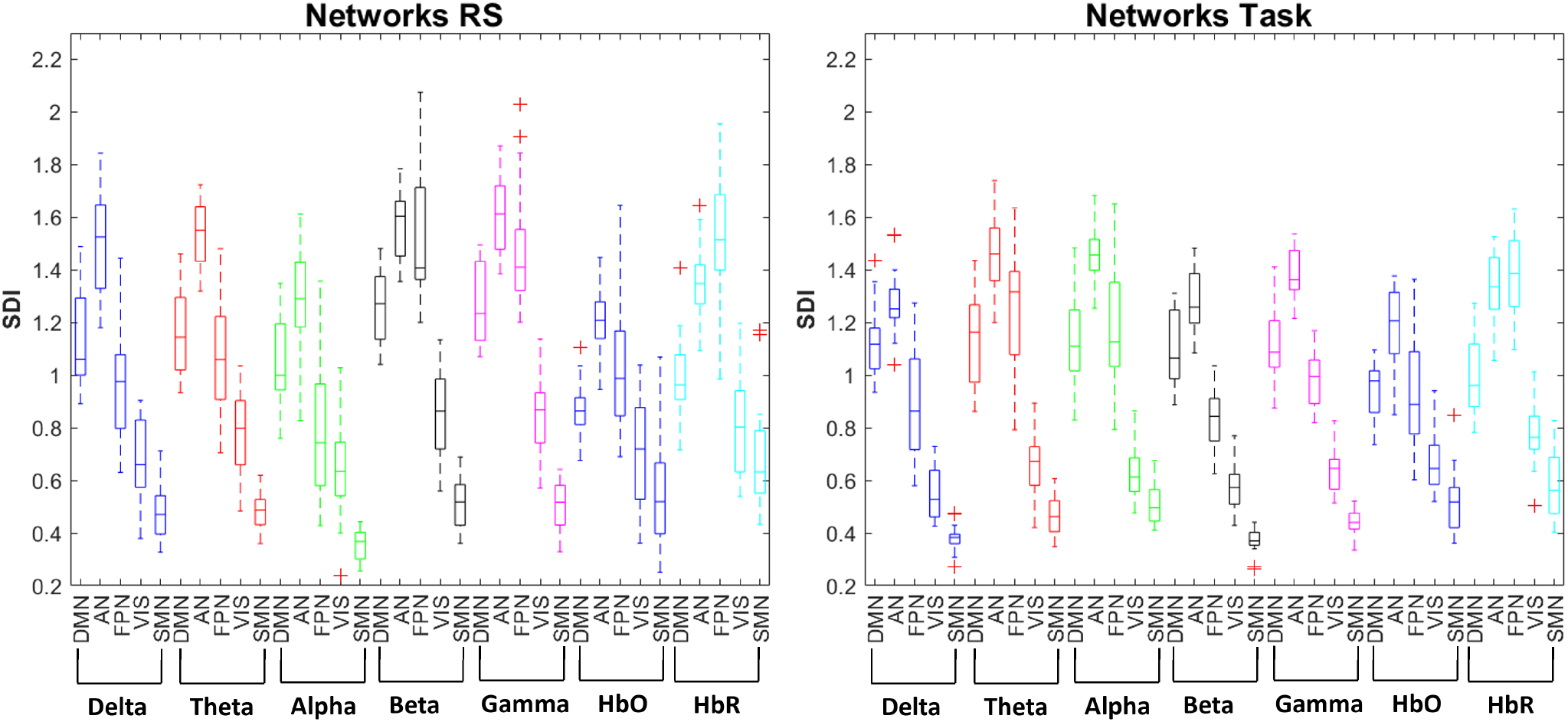
Network structure-function coupling: Boxplot illustrating the distribution of the SDI values across the macro-scale intrinsic networks (DMN, AN, FPN, VIS, and SMN) for each EEG frequency band (Delta, Theta, Alpha, Beta, and Gamma), oxyhemoglobin (HbO), and deoxyhemoglobin (HbR) in resting state (RS) and motor imagery task.

### Network modality-dependent structure-function relationship

Differences emerged when comparing each EEG band with the corresponding HbO and HbR functional networks.

We observed, in RS, greater convergence between EEG and HbO in the primary sensory and motor cortices (VIS and SMN), as well as in the FPN for slower bands (delta, theta), characterized by higher coupling with the structural connectome. A greater divergence between modalities was shown in the transmodal cortex (DMN and AN), characterized by higher decoupling for electrical activity. When compared to faster bands (beta and gamma), greater convergence between the two techniques was only observed in the unimodal cortex (SMN). However, the differences in VIS were marginally significant (*p* = 0.04 for beta and *p* = 0.03 for gamma band) and characterized by higher coupling for hemodynamic activity. Different patterns were highlighted when comparing HbO with the alpha band. A greater convergence between modalities was exhibited in AN (more decoupled) and VIS (more coupled), while divergences were characterized by higher coupling in FPN and SMN and higher decoupling in DMN for EEG.

During the Task, greater convergence between EEG and HbO was shown in the primary sensor and motor cortex (VIS and SMN) within the theta and alpha bands characterized by higher coupling, with greater divergence (higher decoupling for electrical activity) displayed in the transmodal cortex (DMN, AN, and FPN). Within the delta and beta bands, was observed convergence in the AN (higher decoupling) and FPN (higher coupling) and divergence in the DMN, SMN, and VIS, characterized by higher decoupling in the transmodal cortex and higher coupling in the unimodal cortex for electrical activity. Additionally, convergence between modalities was displayed within the gamma band in FPN and VIS (coupled). At the same time, divergence in the DMN, AN, and SMN was characterized by higher decoupling in the transmodal cortex and higher coupling in the unimodal cortex for electrical activity (Fig. 7 (a)). The convergence and divergence patterns between each EEG band and HbR exhibit notable complexity and variation. Briefly, in RS, convergence between EEG and HbR was observed in FPN and VIS within the faster rhythms (beta and gamma), in VIS within the theta band, and in DMN and AN within the alpha band, with greater divergence for all networks within the delta band. During the task period, convergence was exhibited in DMN within the alpha and beta bands, and AN within the delta, beta, and gamma bands (Fig. 7 (b)). The summary of statistical tests is reported in Table 4 of the Supplementary Materials.

**Figure 7.**
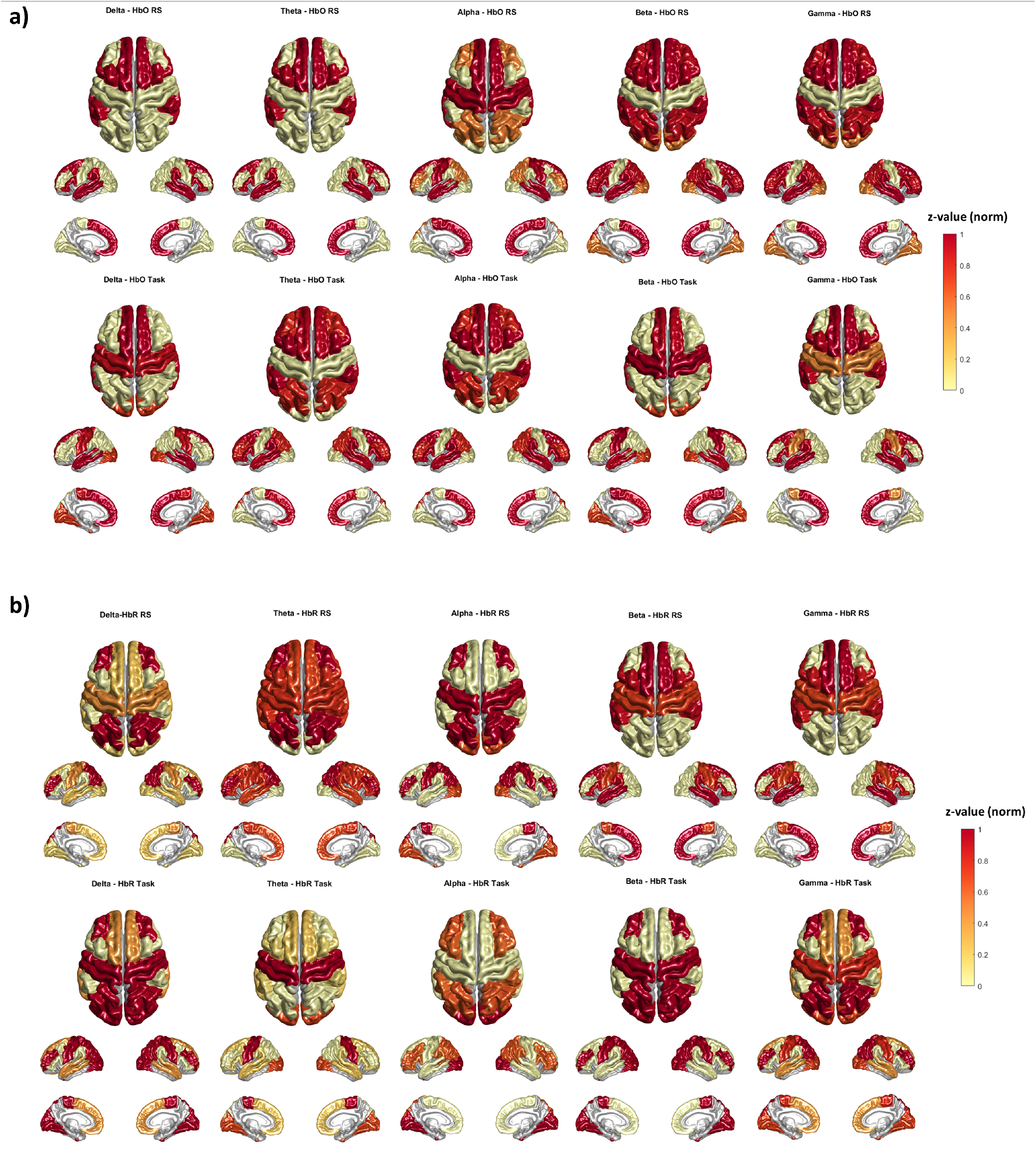
Network structure-function coupling: Brain maps of the differences of the spatial distribution of the network structure-function coupling/decoupling between each EEG frequency band and HbO in **(a)** and between each EEG frequency band and HbR in **(b)** in resting state (RS) and motor imagery task, represented as z-values (norm).

These findings suggest that the spatial organization of oxyhemoglobin structure-function coupling is reminiscent of but distinct from EEG, and dependent on the rhythm being considered and the brain state. This suggests the idea that different EEG rhythms may play distinct roles in modulating cerebral blood flow and oxygenation, thus shaping the regional distribution of hemodynamic responses in the brain.

Indeed, when comparing cross-band EEG with HbO functional networks, differences in structure-function coupling emerged in the DMN and the AN in RS, suggesting divergent interactions between electrical and hemodynamic activity with the underlying structure. During the task, these disparities extended to the SMN, indicating a broader pattern of discordance between EEG and HbO coupling with the structural connectome during the active cognitive engagement (Fig. 8 (a)). Notably, these differences were reflected in greater decoupling for EEG DMN and AN networks in both RS and Task and greater coupling for EEG SMN during the task. No difference was pointed out for the VIS network (Fig. 8 (b)). When comparing cross-band EEG with HbR, only the VIS network in RS and the AN in the task displayed convergence between modalities (Fig. 8 (c)) with a pattern characterized by higher decoupling within the DMN and AN and higher coupling within the FPN, VIS, and SMN for EEG in both RS and task (Fig. 8 (d)). This suggests that the association between structural and functional relationships varies depending on the measurement modality, with electrical activity demonstrating a stronger coupling in sensory-motor networks (SMN), and stronger decoupling in transmodal cortex (DMN, AN) compared to both oxy- and deoxyhemoglobin irrespective of the brain state. In contrast, the coupling pattern in the frontoparietal network (FPN) is condition and modality-dependent. Oxygenated hemodynamic activity exhibits stronger coupling with the underlying structural architecture when the brain is engaged in motor imagery tasks compared to electrical activity, while deoxygenated hemoglobin shows an inverted pattern with lower coupling than electrical activity. The summary of statistical tests is reported in Table 5 of the Supplementary Materials.

**Figure 8.**
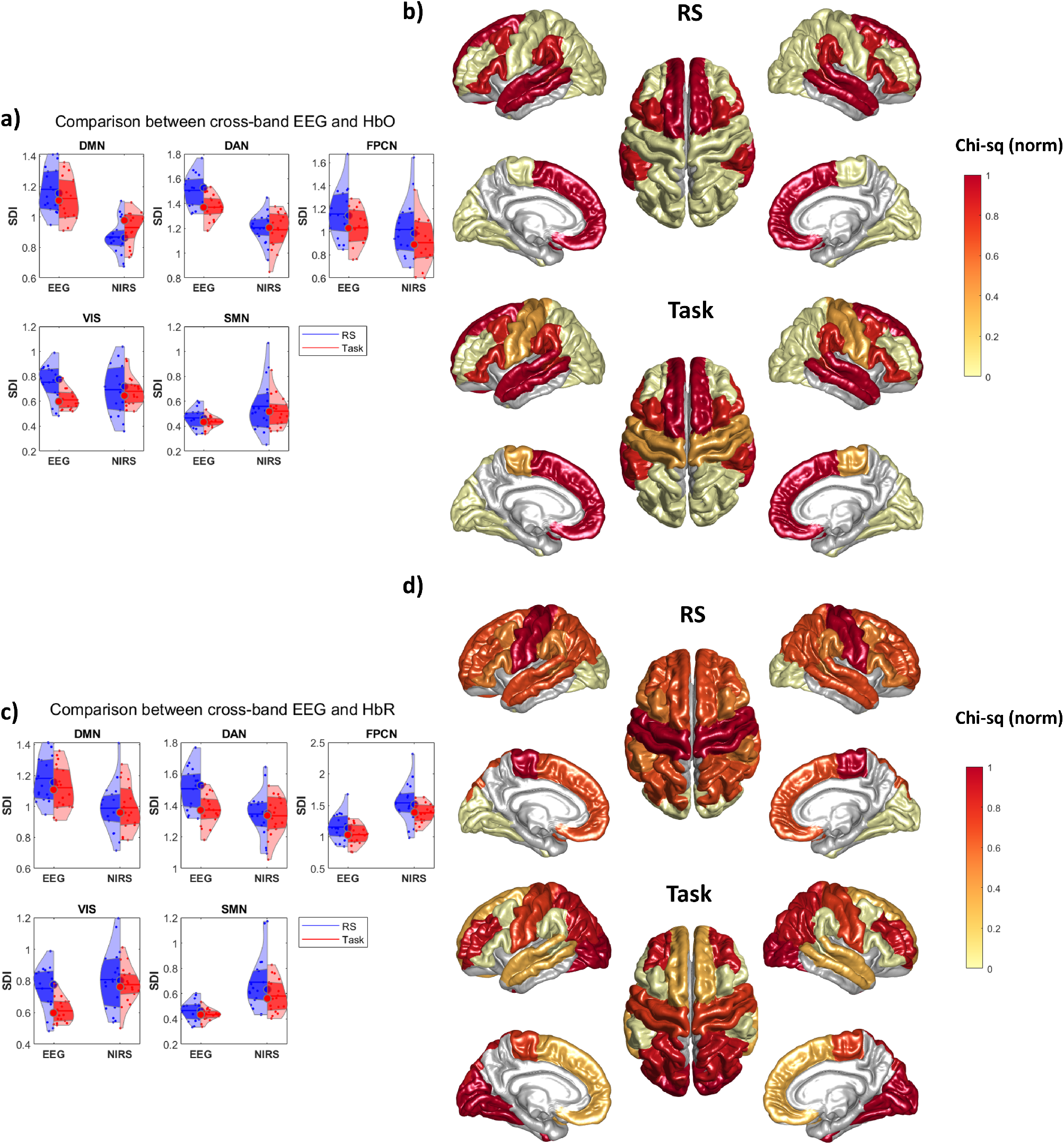
Network structure-function coupling: Violin plot illustrating the differences between **(a)** cross-band EEG and HbO network in RS and task and **(c)** cross-band EEG and HbR network in RS and task. Spatial distribution of the differences between **(b)** cross-band EEG networks and HbO in RS and task and **(d)** cross-band EEG networks and HbR in RS and task, represented as Chi-sq values (norm).

## Discussion

This study assesses the patterns of structure-function coupling throughout the neocortex by analyzing synchronized EEG and fNIRS data during resting state and motor imagery tasks. To our knowledge, this research represents the first investigation into the structure-functional relationship using fNIRS data and comparing it with EEG.

We identified five primary findings. First, the fNIRS structure-function coupling typically resembles slower-frequency EEG structure-function coupling. Second, the structure-function coupling varies across different oscillations between the resting and task periods, alongside the associated hemodynamic response. Third, the structure-function relationship of the hemodynamic activity is specific for oxy- and deoxyhemoglobin in response to neuronal activity. Fourth, local structure-function relationships exhibit heterogeneity but follow a systematic pattern across different modalities and conditions, organized along the sensorimotor association axis. However, discrepancies in specific cortical regions between the two modalities depend on brain state and frequency. Fifth, cross-band representations of neural activity reveal lower correspondence between electrical and hemodynamic activity in the transmodal cortex regardless of brain state while showing specificity for the somatomotor network during a motor imagery task.

At the global level, our results showed that the fNIRS structure-function coupling typically resembles slower-frequency EEG structure-function coupling. However, inter-modality differences, as well as variations among different EEG rhythms, are also evident. As previously reported, we observed that the slower and intermediate neural rhythms (delta, theta, and alpha bands) and hemodynamic activity (oxyhemoglobin) are more coupled with the underlying structure than the faster neural oscillations (beta and gamma) and deoxyhemoglobin [28, 74]. Indeed, the overlap between structural and functional connections depends on the time scale considered, where functional connections estimated from larger time windows strongly overlap with the underlying structural connections. For smaller time windows there can be a structural–functional network discrepancy due to distributed delays between neuronal populations that cause transient phase (de-)synchronization [39, 75].

Moreover, we observed a general trend of increased coupling for both modalities, with no significant difference between the two techniques observed during the task period, in contrast to the RS. This implies that the modalities exhibit more similar patterns of structure-function relationship when the brain is engaged in a specific cognitive task. This phenomenon may indicate the dynamic integration of information across distant brain regions to facilitate task-relevant processes and cognitive functions. It was reported that a higher resemblance to the structural connectome is evident when the functional network is in a highly integrated state [76]. However, increased modular segregation might be manifested, reflecting flexibility away from the structural connectome [59, 76].

We noted variations in structure-function coupling among different oscillations between the resting and task periods, alongside the associated hemodynamic response. Specifically, EEG activity in the alpha band demonstrated a stronger coupling relative to other oscillations at rest, which aligns with the notion that alpha oscillations are linked to an idling or inhibitory state [77]. Contrarily, during the task, the delta and beta bands displayed a more pronounced coupling pattern, indicative of task-specific modulation of neural interactions during motor imagery processing. It could be argued that the specific patterns of structure-function coupling in EEG observed in RS and task conditions might align with the concept of nested oscillations and cross-spectral coupling theory [78]. Indeed, connectome harmonics have been theoretically proposed as a mechanism for macroscopic brain activity, enabling nested functional segregation and integration across multiple spatiotemporal scales [79]. In the case of fNIRS, the hemodynamic activity exhibited an overall relatively stable structure-function relationship across both brain states for both hemoglobin types. Oscillation-based networks are stable over long periods, and as in the case of fNIRS, their organization is largely invariant to changing cognitive demands [1]. However, HbO showed greater alignment with the alpha band in the resting state and the delta band during the task. It was reported that alpha power showed a close fit between the observed and predicted hemodynamic responses as measured by fNIRS [80] and that the modulation of hemodynamics fNIRS response by EEG power was not limited to the alpha band [81, 82, 83, 84]. Moreover, there is some consensus that the best agreement between resting-state fMRI and MEEG/EEG signals is in the alpha and beta bands [85, 86], both in empirical and simulated MEG connectivity based on coupled oscillators with parameters derived from structural networks [46].

Interestingly, HbR showed higher decoupling compared to the alpha rhythm during the RS and to delta and beta oscillations during the task period. These findings are in line with concurrent fNIRS-fMRI studies, which have demonstrated a strong correlation between the spatiotemporal characteristics of HbR and the BOLD signal across various experimental conditions [87, 88, 89, 90, 91, 92].

At the local level, we observed a heterogeneous but systematic structure-function relationship across different modalities and conditions. The cross-modal correspondence between modalities is manifested as consistent patterns of coupling or decoupling within the same ROI, albeit varying in intensity and indicating a predominant coupling of HbO over the different EEG bands than HbR. This pattern revealed greater coupling in the sensory cortex and increased decoupling in the association cortex, following the unimodal to transmodal gradient as previously reported [3, 28, 29]. However, discrepancies in specific cortical regions between the two modalities were also noted, which are condition- and frequency-specific. These discrepancies were pointed out in the right prefrontal and the right posterior parietal cortices. EEG demonstrated higher coupling than fNIRS within slower EEG bands (delta, theta, and alpha) and lower coupling within faster bands (beta and gamma) at rest. Conversely, during the task, we observed a higher coupling in the same regions for the delta, beta, and gamma bands. These findings might agree with the idea that the cortical regions operate at different timescales based on their functional specialization within the cortical hierarchy, suggesting that slower rhythms, primarily localized in deeper layers, are associated with higher-order cognitive processing and integration of sensory information while faster rhythms, primarily localized in superficial layers, are involved in processing more specific or localized information. Previous studies have reported that the correspondence between hemodynamic and electromagnetic connectivity is linked to the underlying cytoarchitectural variation across the cortex, with regions with a higher density of granular cells (unimodal cortex) exhibiting higher cross-modal correspondence and vice versa (transmodal cortex) [28, 93, 94, 95]. Indeed, patterns of neurovascular coupling display a topography with the greatest vascularization density in layer IV [28, 96, 97] which encompasses primary sensory cortices and receives the majority of feedforward input [98, 99, 100].

In light of our results showing variability between modalities which is brain state- and frequency-dependent, and considering the hierarchical model, which delineates separate superficial and deep layers, it is plausible that this architecture could exhibit more intricate spectral characteristics [101, 102]. These may encompass a broader range of temporal dynamics, a complexity that may be challenging to capture with fNIRS, because of its lower temporal resolution, but can be addressed by EEG.

When comparing the intrinsic functional networks between EEG and fNIRS, we found a convergent pattern between modalities characterized by greater structure-function coupling in the visual and somatomotor networks and lower coupling in the default mode and attention network. However, nuances arise in the frontoparietal network, where the structure-function coupling is influenced by frequency bands, hemoglobin types, and brain states. Specifically, greater convergence in the FPN structure-function coupling was observed between HbO and slower frequency bands (delta, theta) in the resting state, while between HbO and delta, beta, and gamma bands during the task. On the contrary, higher convergence was displayed between HbR and beta and gamma bands at rest, while between HbR and theta band during the task.

These findings suggest that the spatial organization of fNIRS structure-function coupling is reminiscent of but distinct from EEG. This resemblance is dependent on the hemoglobin types, the rhythm being considered, and the brain state, implying potentially different modes of intrinsic functional organization in EEG [103, 104], and aligns with existing research on the distinct functional roles of oxyhemoglobin and deoxyhemoglobin [105]. Similarly, specific frequency bands may not directly align with fNIRS functional activity, but different canonical frequency bands and electrophysiological coupling modes can influence hemodynamic responses through neurovascular coupling mechanisms. Indeed, previous research has shown that synchronized oscillations across multiple frequency bands best explain the well-established fMRI functional networks [40].

Therefore, we incorporated the cross-band representations of the electrical networks to gain further insights into the convergence/divergence of EEG and fNIRS structure-function relationship within the intrinsic functional networks. Differences between modalities emerged in the DMN and AN in both the resting and task conditions, confirming a lower correspondence between electrical and hemodynamic activity in the transmodal cortex. These differences extended to the SMN during the task period, indicating a broader pattern of discordance between EEG and HbO structure-function coupling, during an active cognitive engagement. These variations are characterized by greater decoupling in the DMN and AN, alongside increased coupling in the SMN for EEG activity. They may reflect the underlying organization and functional specialization of the human brain [29, 106, 107, 108, 109, 110]. The transmodal cortex integrates information from multiple sensory modalities and higher-order cognitive processes, such as attention, memory, and decision-making. This complex processing may involve distributed neural networks and dynamic interactions between cortical regions, leading to a higher degree of functional segregation and weaker coupling with the underlying structure. This lower coupling might indicate that the transmodal regions are more flexible and adaptable in their functional organization, allowing for dynamic reconfiguration of neural networks to support diverse cognitive tasks [59]. Additionally, this lower coupling may reflect the presence of extensive long-range connections, which facilitate the integration of information across different brain regions. Previous studies provided evidence for the functional role of long-range neuronal coupling in integrating distributed information in the human brain and demonstrate that inter-areal synchronization predicts behavioral performance, illustrating the functional relevance of large-scale coupling for cognitive processing [111, 112, 113]. However, these long-range connections may not exhibit strong coupling with specific structural motifs or local cortical architecture, resulting in a weaker association between functional activity and underlying structural features [114]. Additionally, structural measures were identified, delineating differences across cognitive states, with interhemispheric and local dense intrahemispheric connectivity supporting resting-state function and long-range intrahemispheric connectivity supporting task-driven function [113]. Furthermore, our findings are consistent with prior research, revealing that connectivity within the beta band becomes prominent during task engagement, confirming the significant role of this frequency range in facilitating interaction between distant brain regions [115]. The crucial involvement of the beta band in long-range connectivity aligns with predictions from computational models [116, 117], as well as evidence from invasive recordings in monkeys [118, 119, 120], and clinical studies involving various patient cohorts [121, 122, 123, 124].

Conversely, higher coupling in the somatomotor network could be attributed to its strong and well-defined structural connections that support its specific functions, relying on direct structural connections and leading to stronger coupling with the underlying structural connectome. Therefore, neural activity in the SMN may exhibit rapid and dynamic changes in response to sensory inputs and motor commands and may be more closely tied to the underlying structure, which provides the anatomical framework for signal transmission. Hemodynamic activity measured by fNIRS, which reflects slower changes in blood flow and oxygenation, may not fully capture the rapid dynamics of neural activity in the SMN. Additionally, hemodynamic activity may be influenced by factors such as vascular reactivity and metabolic demands, which could introduce additional variability and lead to weaker coupling with the underlying structural connectivity in the SMN compared to EEG [80, 125].

Our study highlights that structure-function coupling captures characteristic features of the functional organization of the human brain, with these relationships being modality and state-dependent. Specifically, our findings underscore the differential impacts of underlying brain structure on resting and task-driven cognitive states, with clearer frequency effects observed for modalities operating on faster time scales. Additionally, the dynamic nature of the structure-function relationship in the human brain across modalities, brain regions, frequency bands, and brain states emphasizes the need to consider multiple modalities and conditions when investigating mechanisms underlying brain function. These findings set the stage for further exploration into the mechanisms that govern brain function and dysfunction in future research endeavors.

It is essential to acknowledge three key methodological limitations. Firstly, our assessment of structural connectivity relied on diffusion MRI and a constructed group consensus structural connectivity. Although we utilized data from synchronous EEG-fNIRS recordings, our study was constrained by a limited sample size. Future investigations aiming to compare multimodal structure-function relationships could benefit from larger sample sizes and the use of precision imaging techniques that integrate multiple modalities within individual subjects [126, 127]. Secondly, we utilized data with a scan duration of 1 minute for the resting-state condition. While for the task, we captured both electrical and hemodynamic activity at temporal scales able to detect the typical event-related responses (ERPs), for the resting state, this short duration of data, unlike EEG, may not be sufficient for fNIRS due to the slow temporal dynamics of the hemodynamic response to stabilize. However, it has been demonstrated that a 1-minute scan is adequate for obtaining functional connectivity and network metrics stabilization and reproducibility [128]. Future studies aiming to compare multimodal structure-function relationships could explore longer scan durations. Finally, we used a reduced number of regions of interest derived from the Desikan-Killiany atlas due to the limited coverage of fNIRS optodes, which may have led to an incomplete representation of cortical regions and potentially overlooked important functional connections between brain areas. This limitation highlights the need for more comprehensive brain atlases or parcellation schemes that better align with the spatial resolution and coverage of the imaging modalities being used [129, 130]. Future studies could explore alternative high spatial resolution parcellation with higher sensor schemes to provide a more accurate representation of the structure-function relationship across the brain.

## Data Availability

Data used in preparation of this article were obtained from the Open access dataset for simultaneous EEG and NIRS brain-computer interface (BCI)

https://doc.ml.tu-berlin.de/hBCI/contactthanks.php and the Structural connectivity data of the ARCHI database in the Desikan atlas. Human Brain Project Neuroinformatics Platform

https://doi.org/10.25493/91BN-SZ9. The supporting code will be available on GitHub

https://github.com/RosmaryBlancoSano/Structure-Function-Relationship-EEG-and-fNIRS.

## Acknowledgements

The publication was created within the project of the Minister of Science and Higher Education “Support for the activity of Centers of Excellence established in Poland under Horizon 2020” on the basis of the contract number MEiN/2023/DIR/3796. This project has received funding from the European Union’s Horizon 2020 research and innovation programme under grant agreement No 857533. This publication is supported by Sano project carried out within the International Research Agendas programme of the Foundation for Polish Science, co-financed by the European Union under the European Regional Development Fund. Sano Centre for Computational Medicine, Kraków, Poland (https://sano.science/). MGP was supported by the CIBM Center for Biomedical Imaging, a Swiss research center of excellence founded and supported by Lausanne University Hospital (CHUV), University of Lausanne (UNIL), Ecole polytechnique fédérale de Lausanne (EPFL), University of Geneva (UNIGE) and Geneva University Hospitals (HUG).

## Author contributions statement

**R.B**.: Conceptualization, Methodology, Data Analysis, Writing - Original draft preparation, Writing, Visualization. **M.G.P**.: Methodology, Data curation, Writing. **C.K**.: Methodology, Data curation, Writing. **D.V.D.V**.: Supervision. **A.C**.: Supervision. All authors reviewed the manuscript.

