## Supplementary Materials for "Structure-Function Relationship in Electrical and Hemodynamic Brain Networks: Insights from EEG and fNIRS during Rest and Task States"

### ABSTRACT

Identifying relationships between structural and functional networks is crucial for understanding the large-scale organization of the human brain. The potential contribution of emerging techniques like functional near-infrared spectroscopy to investigate the structure-functional relationship has yet to be explored. In our study, we characterize global and local structure-function coupling using source-reconstructed Electroencephalography (EEG) and Functional near-infrared spectroscopy (fNIRS) signals in both resting state and motor imagery tasks, as this relationship during task periods remains underexplored. Employing the mathematical framework of graph signal processing, we investigate how this relationship varies across electrical and hemodynamic networks and different brain states. Results show that fNIRS structure-function coupling resembles slower-frequency EEG coupling at rest, with variations across brain states and oscillations. Locally, the relationship is heterogeneous, with greater coupling in the sensory cortex and increased decoupling in the association cortex, following the unimodal to transmodal gradient. Discrepancies between EEG and fNIRS are noted, particularly in the frontoparietal network. Cross-band representations of neural activity revealed lower correspondence between electrical and hemodynamic activity in the transmodal cortex, irrespective of brain state while showing specificity for the somatomotor network during a motor imagery task. Overall, these findings initiate a multimodal comprehension of structure-function relationship and brain organization when using affordable functional brain imaging.

**Table 1. Global structure-function coupling: Summary of Statistical Tests Comparing EEG and fNIRS Measures Across Different Conditions and Modalities**: Overall comparison between EEG and fNIRS for each condition; Comparison of EEG and fNIRS measures within each modality for each condition; Comparison between conditions for each modality; Comparison between band-limited EEG and fNIRS measures for each condition. P-values (FDR-corrected for multiple comparisons) and z-values are reported for each comparison, indicating the significance of observed differences (in red) or similarities between the measures.

**Global structure-function coupling**

| Overall comparison between modalities | | | |
|---|---|---|---|
| RS | | | |
| Modality_1 | Modality_2 | p-val | z-val |
| *EEG* | *NIRS* | 0.0004 | 3.5490 |
| Task | | | |
| *Modality_1* | *Modality_2* | p-val | z-val |
| *EEG* | *NIRS* | 0.0773 | 1.7663 |
| Comparison between conditions for each modality | | | |
| EEG | p-val | z-val | |
| *RS vs Task delta* | 0.001 | 5.348 | |
| *RS vs Task theta* | 0.376 | 0.886 | |
| *RS vs Task alpha* | 0.001 | -7.734 | |
| *RS vs Task beta* | 0.001 | 12.286 | |
| *RS vs Task gamma* | 0.001 | 9.643 | |
| fNIRS | p-val | z-val | |
| *RS vs Task hbo* | 0.769 | 0.294 | |
| *RS vs Task hbr* | 0.423 | 0.800 | |

| EEG within modality comparison for each condition | | | | | |
|---|---|---|---|---|---|
| RS | | | Task | | |
| | p-val | z-val | | p-val | z-val |
| *delta vs theta* | 0.001 | -6.409 | *delta vs theta* | 0.001 | -9.680 |
| | | 9.594 | *delta vs alpha* | 0.001 | -9.227 |
| *delta vs beta* | 0.001 | -11.789 | *delta vs beta* | ***0.2378*** | -1.181 |
| *delta vs gamma* | 0.001 | -11.827 | *delta vs gamma* | 0.0010 | -8.472 |
| *theta vs alpha* | 0.001 | 12.845 | *theta vs alpha* | ***0.4394*** | 0.773 |
| *theta vs beta* | 0.001 | -9.701 | *theta vs beta* | 0.001 | 9.631 |
| *theta vs gamma* | 0.001 | -9.272 | *theta vs gamma* | *0.0167* | 2.394 |
| *alpha vs beta* | 0.001 | -15.972 | *alpha vs beta* | 0.001 | 11.141 |
| *alpha vs gamma* | 0.001 | -15.451 | *alpha vs gamma* | ***0.0514*** | 1.948 |
| *beta vs gamma* | 0.9656 | 0.043 | *beta vs gamma* | 0.001 | -10.981 |
| fNIRS within modality comparison for each condition | | | | | |
| RS | | | *Task* | | |
| | p-val | z-val | | p-val | z-val |
| *hbo vs hbr* | 0.001 | -7.842 | *hbo vs hbr* | 0.001 | -8.054 |

| Comparison between band-limited EEG and fNIRS for each condition | | | | | |
|---|---|---|---|---|---|
| RS | p-val | z-val | RS | p-val | z-val |
| *delta vs hbo* | 0.001 | 5.725 | *delta vs hbr* | 0.312 | -1.012 |
| *theta vs hbo* | 0.001 | 7.719 | *theta vs hbr* | 0.455 | 0.747 |
| *alpha vs hbo* | 0.445 | 0.765 | *alpha vs hbr* | 0.001 | -5.517 |
| *beta vs hbo* | 0.001 | 10.864 | *beta vs hbr* | 0.001 | 5.466 |
| *gamma vs hbo* | 0.001 | 11.052 | *gamma vs hbr* | 0.001 | 5.652 |
| Task | p-val | z-val | Task | p-val | z-val |
| *delta vs hbo* | 0.248 | 1.155 | *delta vs hbr* | 0.001 | -3.990 |
| *theta vs hbo* | 0.001 | 7.442 | *theta vs hbr* | 0.504 | 0.668 |
| *alpha vs hbo* | 0.001 | 7.196 | *alpha vs hbr* | 0.699 | 0.386 |
| *beta vs hbo* | 0.043 | 2.028 | *beta vs hbr* | 0.001 | -3.889 |
| *gamma vs hbo* | 0.001 | 6.515 | *gamma vs hbr* | 0.454 | -0.749 |

**Table 2.** Mean and std of SDI values across ROIs and subjects for each EEG band and fNIRS chromophore, in RS and task conditions.

| | **Delta** | **Theta** | **Alpha** | **Beta** | **Gamma** | **HbO** | **HbR** |
|---|---|---|---|---|---|---|---|
| **RS** | 1.11 ± 0.12 | 1.16 ± 0.08 | 0.95 ± 0.13 | 1.30 ± 0.09 | 1.30 ± 0.1 | 1.0 ± 0.1 | 1.19 ± 0.09 |
| **Task** | 0.98 ± 0.07 | 1.13 ± 0.11 | 1.11 ± 0.09 | 0.97 ± 0.08 | 1.07 ± 0.07 | 0.96 ± 0.06 | 1.14 ± 0.08 |

**Table 3.** Regions of interest (ROIs) displaying significant disparities between EEG and fNIRS.

| **Number of different ROIs** | | | | | | |
|---|---|---|---|---|---|---|
| | **RS** | **Task** | | | **RS** | **Task** |
| ***Delta vs HbO*** | 24 | 21 | | ***Delta vs HbR*** | 25 | 23 |
| ***Theta vs HbO*** | 28 | 20 | | ***Theta vs HbR*** | 22 | 20 |
| ***Alpha vs HbO*** | 25 | 20 | | ***Alpha vs HbR*** | 27 | 21 |
| ***Beta vs HbO*** | 32 | 22 | | ***Beta vs HbR*** | 20 | 24 |
| ***Gamma vs HbO*** | 32 | 20 | | ***Gamma vs HbR*** | 22 | 22 |

**Table 4.** SDI mean and std for each network, band, and hemoglobin type for the two conditions (RS and Task)

| | **RS** | | | | | | | **Task** | | | | |
|---|---|---|---|---|---|---|---|---|---|---|---|---|
| | **DMN** | **DAN** | **FPN** | **VIS** | **SMN** | | | **DMN** | **DAN** | **FPN** | **VIS** | **SMN** |
| ***Delta*** | 1.14 ± 0.17 | 1.51 ± 0.19 | 0.95 ± 0.21 | 0.67 ± 0.15 | 0.48 ± 0.10 | | ***Delta*** | 1.13 ± 0.13 | 1.27 ± 0.12 | 0.90 ± 0.20 | 0.55 ± 0.09 | 0.38 ± 0.05 |
| ***Theta*** | 1.17 ± 0.16 | 1.54 ± 0.12 | 1.06 ± 0.22 | 0.78 ± 0.16 | 0.49 ± 0.08 | | ***Theta*** | 1.14 ± 0.18 | 1.46 ± 0.15 | 1.25 ± 0.25 | 0.65 ± 0.12 | 0.47 ± 0.08 |
| ***Alpha*** | 1.06 ± 0.16 | 1.3 ± 0.21 | 0.77 ± 0.24 | 0.63 ± 0.18 | 0.36 ± 0.06 | | ***Alpha*** | 1.13 ± 0.18 | 1.45 ± 0.11 | 1.19 ± 0.23 | 0.64 ± 0.11 | 0.51 ± 0.07 |
| ***Beta*** | 1.27 ± 0.14 | 1.58 ± 0.12 | 1.52 ± 0.25 | 0.85 ± 0.15 | 0.51 ± 0.09 | | ***Beta*** | 1.09 ± 0.14 | 1.28 ± 0.12 | 0.84 ± 0.12 | 0.57 ± 0.09 | 0.37 ± 0.04 |
| ***Gamma*** | 1.26 ± 0.14 | 1.6 ± 0.15 | 1.49 ± 0.23 | 0.85 ± 0.14 | 0.52 ± 0.09 | | ***Gamma*** | 1.12 ± 0.13 | 1.39 ± 0.09 | 1.0 ± 0.11 | 0.64 ± 0.08 | 0.44 ± 0.05 |
| ***HbO*** | 0.87 ± 0.11 | 1.2 ± 0.12 | 1.02 ± 0.24 | 0.69 ± 0.19 | 0.56 ± 0.20 | | ***HbO*** | 0.93 ± 0.10 | 1.19 ± 0.14 | 0.91 ± 0.20 | 0.68 ± 0.12 | 0.52 ± 0.12 |
| ***HbR*** | 0.99 ± 0.16 | 1.35 ± 0.14 | 1.54 ± 0.31 | 0.81 ± 0.19 | 0.69 ± 0.21 | | ***HbR*** | 0.99 ± 0.15 | 1.33 ± 0.12 | 1.38 ± 0.15 | 0.78 ± 0.12 | 0.58 ± 0.13 |

**Table 5. Network structure-function coupling: Summary of Statistical Tests Comparing band-limited EEG and fNIRS Measures Across Different Conditions**. P-values and z-values are reported for each comparison, indicating the significance of observed differences (in red) or similarities between the measures.

| | RS | | | | | | | | | |
|---|---|---|---|---|---|---|---|---|---|---|
| **Network** | **p_delta_HbO** | **p_theta_HbO** | **p_alpha_HbO** | **p_beta_HbO** | **p_gamma_HbO** | **p_delta_HbR** | **p_theta_HbR** | **p_alpha_HbR** | **p_beta_HbR** | **p_gamma_HbR** |
| **DMN** | 0.0004 | 0.0002 | 0.0007 | 0.0002 | 0.0002 | 0.0108 | 0.0084 | 0.2145 | 0.0003 | 0.0003 |
| **AN** | 0.0005 | 0.0004 | 0.157 | 0.0003 | 0.0002 | 0.0279 | 0.0038 | 0.6475 | 0.001 | 0.0014 |
| **FPN** | 0.3061 | 0.3271 | 0.0249 | 0.0006 | 0.0007 | 0.0002 | 0.0002 | 0.0002 | 0.9133 | 0.7112 |
| **VIS** | 0.5862 | 0.3271 | 0.3061 | 0.0429 | 0.0347 | 0.0156 | 0.7771 | 0.0096 | 0.3958 | 0.4204 |
| **SMN** | 0.3491 | 0.3958 | 0.0009 | 0.7439 | 0.5566 | 0.0043 | 0.0043 | 0.0002 | 0.0096 | 0.0084 |
| | **RS** | | | | | | | | | |
| | **z-stat_delta_HbO** | **z-stat_theta_HbO** | **z-stat_alpha_HbO** | **z-stat_beta_HbO** | **z-stat_gamma_HbO** | **z-stat_delta_HbR** | **z-stat_theta_HbR** | **z-stat_alpha_HbR** | **z-stat_beta_HbR** | **z-stat_gamma_HbR** |
| **DMN** | 3.5494 | 3.68 | 3.3752 | 3.7236 | 3.7236 | 2.5477 | 2.6348 | 1.2412 | 3.5929 | 3.6365 |
| **AN** | 3.4623 | 3.5494 | 1.4154 | 3.6365 | 3.68 | 2.1993 | 2.8961 | -0.4573 | 3.2881 | 3.201 |
| **FPN** | -1.0234 | 0.9799 | -2.2428 | 3.4187 | 3.3752 | -3.68 | -3.68 | -3.7236 | -0.1089 | -0.3702 |
| **VIS** | -0.5444 | 0.9799 | -1.0234 | 2.0251 | 2.1122 | -2.417 | -0.2831 | -2.5912 | 0.8492 | 0.8057 |
| **SMN** | -0.9363 | -0.8492 | -3.3316 | -0.3266 | -0.5879 | -2.8525 | -2.8525 | -3.68 | -2.5912 | -2.6348 |
| | **Task** | | | | | | | | | |
| **Network** | **p_delta_HbO** | **p_theta_HbO** | **p_alpha_HbO** | **p_beta_HbO** | **p_gamma_HbO** | **p_delta_HbR** | **p_theta_HbR** | **p_alpha_HbR** | **p_beta_HbR** | **p_gamma_HbR** |
| **DMN** | 0.0002 | 0.0006 | 0.0006 | 0.0009 | 0.0005 | 0.0198 | 0.0347 | 0.0778 | 0.1221 | 0.0386 |
| **AN** | 0.1119 | 0.0004 | 0.0005 | 0.1446 | 0.0009 | 0.1841 | 0.0429 | 0.0108 | 0.3061 | 0.3491 |
| **FPN** | 0.8789 | 0.0029 | 0.005 | 0.4204 | 0.1841 | 0.0002 | 0.0582 | 0.0084 | 0.0002 | 0.0002 |
| **VIS** | 0.0057 | 0.5566 | 0.5277 | 0.0074 | 0.4204 | 0.0002 | 0.0108 | 0.0016 | 0.0002 | 0.0038 |
| **SMN** | 0.0007 | 0.157 | 0.9826 | 0.0005 | 0.0386 | 0.0002 | 0.0043 | 0.0854 | 0.0002 | 0.0016 |
| | **Task** | | | | | | | | | |
| | **z-stat_delta_HbO** | **z-stat_theta_HbO** | **z-stat_alpha_HbO** | **z-stat_beta_HbO** | **z-stat_gamma_HbO** | **z-stat_delta_HbR** | **z-stat_theta_HbR** | **z-stat_alpha_HbR** | **z-stat_beta_HbR** | **z-stat_gamma_HbR** |
| **DMN** | 3.7236 | 3.4187 | 3.4187 | 3.3316 | 3.4623 | 2.3299 | 2.1122 | 1.7638 | 1.546 | 2.0686 |
| **AN** | 1.5896 | 3.5494 | 3.5058 | 1.4589 | 3.3316 | -1.3283 | 2.0251 | 2.5477 | -1.0234 | 0.9363 |
| **FPN** | -0.1524 | 2.9832 | 2.809 | -0.8057 | 1.3283 | -3.68 | -1.8944 | -2.6348 | -3.7236 | -3.7236 |
| **VIS** | -2.7654 | -0.5879 | -0.6315 | -2.6783 | -0.8057 | -3.7236 | -2.5477 | -3.1574 | -3.68 | -2.8961 |
| **SMN** | -3.3752 | -1.4154 | -0.0218 | -3.4623 | -2.0686 | -3.7236 | -2.8525 | -1.7202 | -3.7236 | -3.1574 |

**Table 6. Network structure-function coupling: Summary of Statistical Tests Comparing cross-band EEG and fNIRS Measures Across Different Conditions**. P-values and z-values are reported for each comparison, indicating the significance of observed differences (in red) or similarities between the measures.

| RS | | | | | | | | | RS | | | | | | | |
|---|---|---|---|---|---|---|---|---|---|---|---|---|---|---|---|---|
| **Group1** | **Group2** | **Network** | **CI_95L** | **dm** | **CI_95H** | **p_value** | **Chi_sq** | | **Group1** | **Group2** | **Network** | **CI_95L** | **dm** | **CI_95H** | **p_value** | **Chi_sq** |
| ***EEG*** | ***HbO*** | DMN | 30.883 | 46.733 | 62.583 | 0.001 | 33.395 | | ***EEG*** | ***HbR*** | DMN | 15.950 | 31.800 | 47.650 | 0.001 | 15.462 |
| ***EEG*** | ***HbO*** | DAN | 27.750 | 43.600 | 59.450 | 0.001 | 29.067 | | ***EEG*** | ***HbR*** | DAN | 11.550 | 27.400 | 43.250 | 0.001 | 11.480 |
| ***EEG*** | ***HbO*** | FPN | -3.517 | 12.333 | 28.183 | 0.162 | 2.326 | | ***EEG*** | ***HbR*** | FPN | -47.317 | -31.467 | -15.617 | 0.001 | 15.140 |
| ***EEG*** | ***HbO*** | VIS | -6.983 | 8.867 | 24.717 | 0.273 | 1.202 | | ***EEG*** | ***HbR*** | VIS | -22.783 | -6.933 | 8.917 | 0.391 | 0.735 |
| ***EEG*** | ***HbO*** | SMN | -29.183 | -13.333 | 2.517 | 0.159 | 2.718 | | ***EEG*** | ***HbR*** | SMN | -54.117 | -38.267 | -22.417 | 0.001 | 22.390 |
| **Task** | | | | | | | | | **Task** | | | | | | | |
| **Group1** | **Group2** | **Network** | **CI_95L** | **dm** | **CI_95H** | **p_value** | **Chi_sq** | | **Group1** | **Group2** | **Network** | **CI_95L** | **dm** | **CI_95H** | **p_value** | **Chi_sq** |
| ***EEG*** | ***HbO*** | DMN | 21.350 | 37.200 | 53.050 | 0.001 | 21.160 | | ***EEG*** | ***HbR*** | DMN | 8.483 | 24.333 | 40.183 | 0.003 | 9.054 |
| ***EEG*** | ***HbO*** | DAN | 17.883 | 33.733 | 49.583 | 0.001 | 17.400 | | ***EEG*** | ***HbR*** | DAN | -8.783 | 7.067 | 22.917 | 0.382 | 0.764 |
| ***EEG*** | ***HbO*** | FPN | -0.117 | 15.733 | 31.583 | 0.052 | 3.785 | | ***EEG*** | ***HbR*** | FPN | -55.917 | -40.067 | -24.217 | 0.001 | 24.546 |
| ***EEG*** | ***HbO*** | VIS | -32.450 | -16.600 | -0.750 | 0.050 | 4.213 | | ***EEG*** | ***HbR*** | VIS | -53.583 | -37.733 | -21.883 | 0.001 | 21.771 |
| ***EEG*** | ***HbO*** | SMN | -38.117 | -22.267 | -6.417 | 0.006 | 7.581 | | ***EEG*** | ***HbR*** | SMN | -52.250 | -36.400 | -20.550 | 0.001 | 20.259 |
